# In vitro propagation of Okra accessions (Abelmoschus esculentus l. Moench) through Nodal Cuttings

**DOI:** 10.64898/2026.10.28.762680

**Authors:** Samuel Arega Asfaw, Meseret Tesema, Zerihun Demerew, Sualih Gobeze, Wondye Ayaliew Shiferaw

## Abstract

Okra (<u>Abelmoschus esculentus</u> l. Moench) is an important multi-purpose vegetable crop and conventionally propagated by seed. Therefore, this study was set up to develop a regeneration protocol for three Okra accessions through nodal cuttings. The treatments were laid out in factorial arrangement of completely randomized design (CRD) design. Days to shoot and root initiation; percent of root and shoot initiation; number of shoot, leaf and root; length of shoot and root and survival rate of acclimatized plantlets were recorded for mean comparison. To induce shoot from nodal cuttings, MS medium supplemented with three levels (0.5, 1, and 1.5 mg/l) of 6-benzylaminopurinem (BAP) in combination with three levels (0.5, 1, and 1.5mg/l) of á-naphthalene acetic acid (NAA) were used. The maximum mean number of shoots was recorded on MS medium supplemented with 1.5 mg/l BAP + 2 mg/l Silver nitrate (AgNO_3_) for the three accessions. The maximum mean number of roots were recorded on media supplemented with 2mg/l Indole-3-butyric acid (IBA) and at accession Mandura. The tallest length of root was achieved on media without growth regulator and at accession Mandura. The maximum survival rate was recorded on accession Mandura followed by accession Akobo and Diga Leka after a month of acclimatization. Finally, in this study an efficient regeneration protocol from nodal cutting experiments was developed for large-scale multiplication, propagation and conservation of Okra.

## INTRODUCTION

### Back ground and Justification of the Study

Okra (*Abelmoschus esculentus* L. Moench) is an important multi-purpose vegetable crop cultivated in sub-tropical and tropical areas throughout the world (Dhande *et al*., 2012). Okra is a member of the Malvaceae family and is mostly known as lady’s finger (Schippers, 2000). India is the leading country followed by Nigeria, Sudan, Mali and Pakistan with the production of 5.5 million tons and covers an areas of 0.48 million hectares in the world (FAOSTAT, 2016). In Ethiopia, there is no complete record of the production area and production of okra, but okra is a traditional crop in southwestern, western, and northwestern Ethiopia (Yonas *et al*., 2014).

Okra plays an important role in human diet by providing carbohydrates, fats, proteins, vitamins, and minerals (El-Kader *et al*., 2010). It grown mainly for its leaves and young pods, which frequently eaten as green vegetables. Therefore, okra consumption plays an important role in human nutrition. Okra mucilage is suitable for industrial and medicinal applications (Akinyele & Temikotan, 2007). Industrially, okra mucilage used for glace paper production and confectionery. Okra has medical applications as a plasma replacement or blood volume expander (Lengsfeld *et al*., 2004).

Okra is a crop that has the potential to improve food security, reduce malnutrition, and alleviate poverty in tropical and subtropical regions of the world. It is considered a prized vegetable owing to its high nutritive value (Dabire-Binso *et al*., 2009). Owing to its medicinal and nutritional value, okra is widely cultivated in tropical and subtropical regions (Roy *et al*., 2014).

Okra attacked by various pathogens, such as bacteria, viruses, and root-knot nematodes. These pathogens cause biotic stress in plants (Kucharek, 2004). Micropropagation techniques (plant tissue culture) offer an opportunity for the large-scale production of uniform disease-free planting material in a relatively short period of time and are independent of the season (Trigiano & Gray, 2016). *In vitro* derived plants are frequently more vigorous and of superior quality than those produced by *in vivo* methods. *In vitro* systems have been reported as an effective tool for obtaining genetically uniform plants, which can be a source of less variable pharmaceutical preparations (Santarém & Astarita, 2003). Furthermore, plant tissue and cell cultures are important tools that allow extensive manipulation of the biosynthesis of secondary compounds and yield a higher productivity than that of intact plants (Kirakosyan *et al*., 2001).

The ability to regenerate okra plants *in vitro* would permit rapid propagation of this important horticultural crop. Very few reports are available regarding the production of somatic embryos from callused okra hypocotyls. The morphogenic responses of the hypocotyl, cotyledon, cotyledonary node, and primary leaf explants in different media combinations (Blackmon *et al*., 1981). Direct and indirect plant regeneration was achieved from hypocotyl explants (Mangat & Roy, 1986) and successful micropropagation of *Abelmoschus* plant*s* was achieved (Haider *et al*., 1993). Therefore, it is important to apply tissue and organ culture techniques to explore the possibility of okra regeneration. The production of true-to-type planting materials can rapidly achieved through *in vitro* propagation, also termed micropropagation, because of the small size of the plant material used. This propagation method can be performed regardless of the growing season and could therefore enable year-round availability of planting materials and expansion and intensification of production. However, *in vitro* regeneration of plants is influenced by many factors, such as the choice of explant, age of explant, proper choice of culture media, plant growth regulators, genotype, source of carbohydrate, gelling agent, and other physical factors, including light, temperature, and humidity (Feyissa *et al*., 2005).

Among the factors influencing *in vitro* regeneration, plant growth regulators (PGRs) in the media and genotypes were the most important. PGRs (e.g., auxins and cytokinins) play an important role in organogenesis because their regimes can be used to manipulate the morphogenic response of plants under *in vitro* cultures (Arya *et al*., 2009). Both callus induction and plant regeneration from explants require the presence of appropriate concentrations and combinations of plant growth regulators in the culture media (Kalimuthu *et al*., 2007).

The capacity for *in vitro* regeneration varies considerably between plant species. Some families and genera, such as Solanaceae (*Nicotiana, Petunia* and *Datura*), Cruciferae (*Brassica* and *Arabidopsis*), Gesneriaceae (*Achimenes* and *Streptocarpus*), Asteraceae (*Chichorium* and *Chrysanthemum*), and Liliaceae (*Lilium* and *Allium*) can have a high regeneration capacity. In contrast, regeneration in families such as Malvaceae (*Gossypium*) and Chenopodiaceae (*Beta*) is challenging, suggesting that the mechanisms required for *in vitro* regeneration are insufficient or absent (Yildiz, 2012).

In Ethiopia, the development and application of tissue culture techniques for the regeneration of okra is at an early stage. Prior to this study, no documented reports were available on the micropropagation of Ethiopian okra accessions for mass propagation. Considering the importance of the crop and its propagation limitation, the present study was designed to develop *in vitro* protocol for the regeneration of three Ethiopian Okra accessions using nodal cuttings.

## MATERIALS AND METHODS

### Description of the Experimental Site

The present study was carried out in the tissue culture laboratory of Hawassa University, College of Agriculture, during 2018/2019. Hawassa is located in the Sidama Region, Ethiopia, at an altitude of 1700 m a. s.l. (NMA, 2012)

### Experimental Plant Material

The seeds of three okra accessions, Akobo, Digaleka, and Mandura, were collected from the Ethiopian Biodiversity Institute, Addis Ababa, Ethiopia, and used for tissue culture experiments (Table 1).

**Table 1.** Source and growing altitude of planting material of Okra accessions used in the present study.

| **Region** | **Place of collection** | **Altitude (masl)** | **Collectors (Seed source)** | **Accession number (code)** |
| --- | --- | --- | --- | --- |
| Gambella | Akobo | 740 | EBI | 240587 |
| Oromiya | Diga leka | 1200 | EBI | 92203 |
| Benishangulgumuz | Mandura | 1200 | EBI | 240204 |

**Note:** EBI= Ethiopian Biodiversity Institute, masl= meter above sea level

### Treatments and Experimental Design

In the present study, there were four experimental stages (shoot regeneration, shoot multiplication, root induction, and acclimatization). For the first three stages of the experiment, MS media as indicated in (Appendix table 1) were used for regeneration protocol development (Murashige & Skoog, 1962) and for the last stage (acclimatization stage), the soil, compost, and sand composition were at a ratio of 2:1:1 respectively was used. In each developmental stage, three different accessions of okra as indicated in Table 1 were used as factorial combined treatments. Treatments at all stages were established as follows:

### Stage I: Shoot Initiation

To initiate shoots, nodal cuttings of (1-2 cm) from the three Okra accessions were used. For this stage, a combination of three levels of cytokinin BAP (0.5, 1.0, and 1.5 mg/l), three levels of auxin NAA (0.5, 1 and 1.5 mg/l), and three accessions of Okra were used. This resulted in 27 treatments, which were replicated five times. Culture medium without growth hormones was used as a control.

### Stage II: Shoot Multiplication

For multiplication of shoots, BAP at different concentrations (0.1, 0.25, 0.5, 1.0 and 1.5 mg/l) and AgNO_3_ at a concentration of (2 mg/l) were used for the multiplication of three accessions of Okra. At this stage, 15 treatments were arranged in three replicates. MS medium without growth regulators and AgNO_3_ were used as controls.

### Stage III: Root Induction

At this stage, auxin (IBA) at concentrations of (0.25, 0.5, 1.0, 1.5 and 2.00 mg/l) was used for three accessions of Okra. At this stage, 15 treatments were arranged in three replicates. MS medium without a growth regulator used as the control.

### General Procedure for *In Vitro* Regeneration of Okra Preparation of Media Stock Solution

MS medium is widely used as a basal medium for the micropropagation of plant species. The stock solutions of these media consisted of basic macro-and micronutrients, and organic supplements (Appendix 1). Stock solutions were prepared according to the recommended amounts of the chemicals, weighed using an electronic sensitive balance, and dissolved in distilled water. Stock solutions were poured into labeled bottles and stored in a refrigerator at 4 ^0^C. The nutrient media were prepared by taking the recommended amounts of MS stock solutions (100 ml/1 macro, 1 ml/l micro, and 1 ml/l vitamins) supplemented with 3% (w/v) sucrose and 0.8% (w/v) agar as solidifying agent. A solidifying agent (agar) was added to the medium prior to autoclaving after adjusting the pH to 5.8 with the help of 1N NaOH or HCL. All media were autoclaved at 121 °C for 20 min. Finally, the autoclaved media were retained in the transfer room for a maximum of two days prior to use.

### Preparation of Plant Growth Regulators Stock Solution

MS basal medium was used for the current study, which is commonly used for organogenesis and regeneration of plants in tissue culture (Murashige & Skoog, 1962). The PGRs used in this experiment were BAP, NAA, and IBA. All PGR stock solutions were prepared by weighing the required amount of the chemical using an electronic sensitive balance and dissolving it in sterilized distilled water at a ratio of 1 mg/ml. To begin the dissolving process, the powdered crystals of the PGRs were first weighed and dissolved in 3-4 drops of 1N NaOH and 1N HCL based on the type of PGRs (NaOH for auxins and HCL for cytokinins). Thereafter, each stock solution adjusted to the desired volume in a volumetric flask using sterilized distilled water. Finally, the PGR stock solutions were poured in to properly labeled media bottles and stored in a refrigerator at the temperature of 4 °C for short-term use.

### Preparation of Mother Stock

The seeds of three Okra accessions were germinated in plastic pots containing soil, sand, and manure in a 2:1:1 ratio and kept as mother plants. The plants were maintained in a net house for approximately one month and watered daily. After a month of growth, very young, healthy, and vigorous parts of the plant (nodes) were collected and used as a source of explants to develop regeneration protocols for the three accessions.

### Sterilization of Planting Materials

The explants obtained from the mother plant, which was cultivated in a net house. The explants/nodes taken from the mother plant sterilized before performing further experiments. The explants were gently washed under running tap water and then with distilled water to remove dust particles. The washed with disinfectant dettol followed by washed three times rising with distilled water then sterilized with 70% (v/v) ethanol for 3 min, followed again by three rinses (3 min each) with sterilized distilled water. The explant surface was disinfected with sodium hypochlorite (commercial bleach) containing 3-4 drops of Tween-20 per 100 ml for 15 min, and then rinsed five times (3 min each) with sterilized distilled water.

### Culturing Conditions of the Explants

All stages of the present study (shoot initiation, shoot multiplication, and root induction) cultured under an 8 h dark period and 16 h photoperiod in a growth room. Artificial light was provided using parallel cool white fluorescent tubes installed above the culture. The light intensity was regulated to 1500-2000 lux and the growth room temperature was adjusted to 25±2 °C with relative humidity (RH) of 65-70%.

### Acclimatization

Well-rooted *in vitro* propagated explants were removed from the culture medium without damaging the root, washed thoroughly in sterilized water to remove all the residues of the medium, including agar, planted in labeled pots containing soil, compost, and sand in a ratio of 2:1:1, and then watered every day in the greenhouse for four days. To control the microclimate, such as humidity, it was covered with plastic bags, random holes were made for air circulation, and the underside of the pots was drilled for drainage. The acclimatized plantlets were then transferred to a net house for further hardening.

### Data Collected

The following plant growth and morphological data were collected:

✓ Days to shoot initiation
✓ Percent of shoot initiated
✓ Mean number of shoots, leaves
✓ Mean length of shoots
✓ Days to root initiation
✓ Percent of root initiated
✓ Mean number of root
✓ Mean length of root
✓ Survival rate

### Analysis of Data

The treatment effects in all experiments were determined using analysis of variance (ANOVA), and significant differences among treatments were determined by Fisher’s least significant difference (LSD) (P<0.05) using the SAS software package version 9.0 (SAS, 2002). For all the data analysis, probability of 5% was considered for statistically significant.

## RESULTS AND DISCUSSION

The present study is generally involved in the development of *in vitro* regeneration protocol for three okra accessions collected from different parts of Ethiopia. The experiment was performed using nodal cutting as an explant for organ regeneration. *In vitro* regeneration of plants involves direct and indirect organ regeneration. In direct regeneration, shoots or roots are directly induced from tissue explants, whereas indirect organogenesis involves callus formation as an intermediate prior to shoot or root induction. In the present study, direct organogenesis performed using nodal cuttings.

### Shoot Initiation from Nodal Explants

In the present study, shoots were initiated from nodal explants of three accessions of okra on a medium containing cytokinin (BAP) and auxin (NAA) in combination (Figure 1). Days to shoot initiation and the percentage of shoot initiation recorded for the initiated nodal explant.

**Figure 1.**
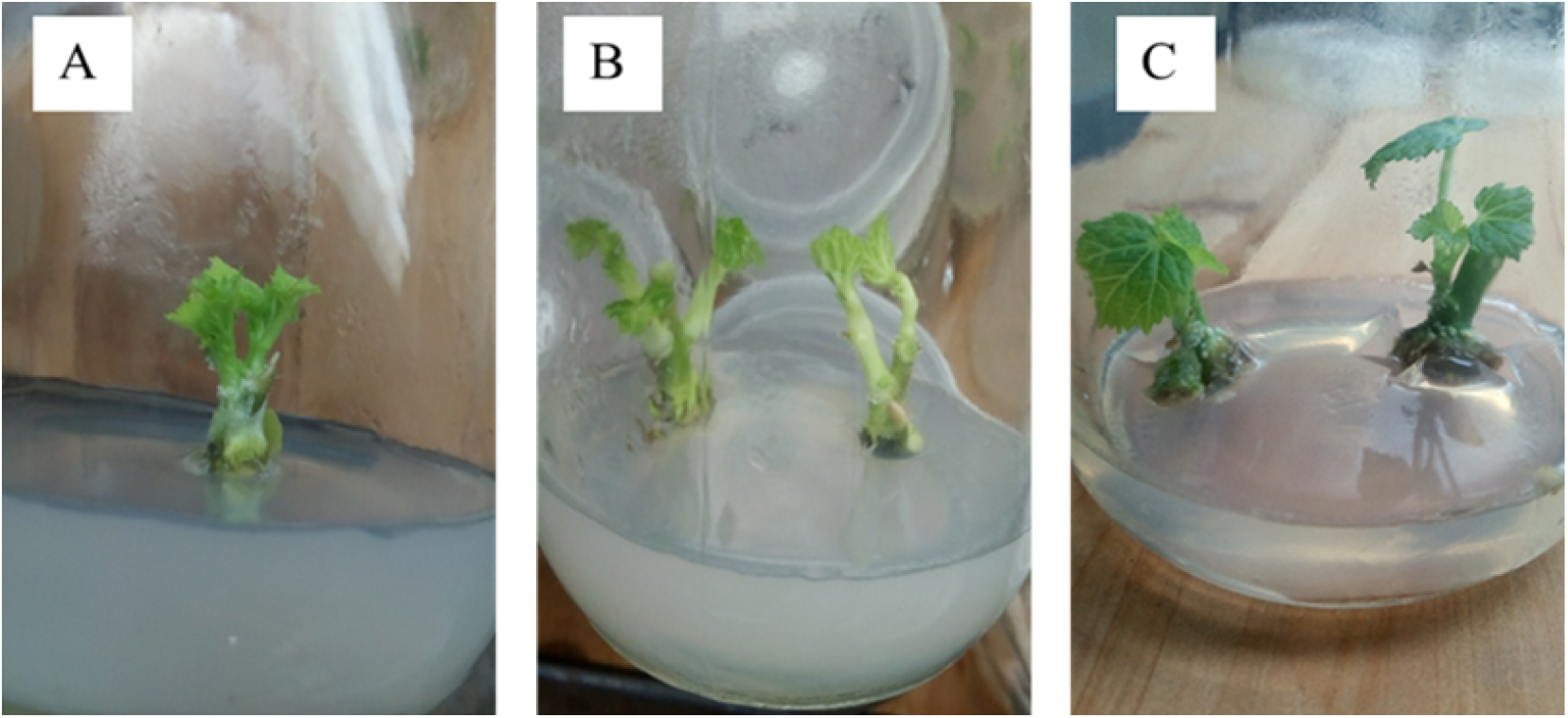
Initiated shoots of Okra accessions after 10 days of culture from nodal explant regenerated on media containing: A) 0.5mg/l BAP with 0.5mg/l NAA of Mandura accession; B) 1.5mg/l BAP with 0.5mg/l NAA of Akobo accession; C) 1mg/l BAP with 0.5mg/l NAA of Diga leka accession.

### Days to Shoot Initiation

Analysis of variance revealed that the interaction of PGRs and accessions had no significant effect (P>0.05) on days to shoot initiation (Table A2). The number of days required for shoot initiation was significantly (P<0.01) influenced by the main effect of the growth regulator concentration and combinations. Considerable variation was observed in the days to shoot initiation in explants cultured on different concentrations of plant growth regulators for all three accessions (Table 2).

**Table 2.** Effect of PGRs (BAP and NAA) combinations on days to shoot initiation and percentage of initiation of *in vitro* nodal culture of three Okra Accessions.

| PGRs concentration (mg/l) |  | Days to shoot initiation | Percent of shoot initiated |
| --- | --- | --- | --- |
| NAA | BAP |  |  |
| 0 | 0 | 11.33±0.14<sup>a</sup> | 0.48±0.04<sup>e</sup> |
| 0.5 | 0.5 | 7.6 ±0.14<sup>e</sup> | 0.77±0.04<sup>bc</sup> |
| 1 | 0.5 | 8.93±0.14<sup>c</sup> | 0.72±0.04<sup>c</sup> |
| 1.5 | 0.5 | 9.4±0.14<sup>b</sup> | 0.53±0.04<sup>de</sup> |
| 0.5 | 1 | 5.93±0.14<sup>g</sup> | 0.85±0.04<sup>a</sup> |
| 1 | 1 | 7.67 ±0.14<sup>e</sup> | 0.73±0.04<sup>c</sup> |
| 1.5 | 1 | 8.4±0.14<sup>d</sup> | 0.58±0.04<sup>d</sup> |
| 0.5 | 1.5 | 6.73±0.14<sup>f</sup> | 0.83±0.04<sup>ab</sup> |
| 1 | 1.5 | 9±0.14<sup>c</sup> | 0.72±0.04<sup>c</sup> |
| 1.5 | 1.5 | 9.47 ±0.14<sup>b</sup> | 0.58±0.04<sup>d</sup> |
| LSD (5%) |  | 0.37 | 0.08 |
| CV (%) |  | 6.05 | 15.45 |

CV, coefficient of variation; LSD= Least Significant Difference; SE= Standard Error; PGR= Plant Growth Regulator; BAP, 6-benzylaminopurine; NAA, $\alpha$ -naphthalene Acetic Acid. Means with different letters in a column are statistically significant at $p < 0.05$ . Values are presented as the mean ± SE.

The minimum number of days to shoot initiation was recorded from explants cultured on MS medium supplemented with a combination of 1 mg/l BAP and 0.5 mg/l NAA followed by 1.5 mg/l BAP and 0.5 mg/l NAA for all accessions. Whilst, the maximum mean days to shoot initiation was recorded from explants cultured on the media without growth regulators followed by 1.5 mg/l BAP with 1.5 mg/l NAA and 0.5 mg/l BAP and 1.5 mg/l NAA for three of Okra accessions.

The results of the present study revealed that the number of days to initiate shoot increased with increasing in NAA concentration of NAA in the treatments by 29% from 0.5 mg/l of NAA to 1.5mg/l of NAA. A high concentration of BAP, in combination with a low concentration of NAA, resulted in the minimum number of days to shoot initiation. This indicates that increasing the concentration of cytokinin (BAP) is important for shoot initiation. High concentration of auxins is generally inhibitory to morphogenesis and substitution of these reagents with an appropriate auxin-cytokinin is essential to obtain proper shoot primordial (Kalimuthu *et al*., 2007). Hormonal balance is believed to be a key factor in regulating explant morphogenesis in explants (Dhande *et al*., 2012). This proves that the cross-talk between auxin and cytokinin during *in vitro* organogenesis is very important for achieving easy and stable establishment of explants. Studies on the classical chemical regulation of growth and organ formation in *in vitro* plant tissues have indicated that excess cytokinin promotes shoot formation, whereas a low ratio of cytokinin to auxins promotes callus induction (Murashige & Skoog, 1962). Hence, in the present study, the increasing number of days to shoot formation with increasing concentrations of NAA may be the result of the inhibitory effects of auxin.

### Percentage of Shoot Initiation

Analysis of variance showed that there was no significant difference (P>0.05) among the interaction effects. Similarly, there were no significant differences (P>0.05) among the three accessions. However, there was a highly significant difference (P<0.01) between plant growth regulator concentrations and combinations (BAP and NAA) in the percent of shoot initiation (Table A2).

The highest percentage to shoot initiation for all accessions was recorded for explants cultured on MS medium supplemented with combination of 1 mg/l BAP and 0.5 mg/l NAA followed by 1.5 mg/l BAP with 0.5 mg/l NAA (Table 2). Whereas, the lowest percentage to shoot initiation was recorded for explants cultured on the media without growth hormones followed by 0.5 mg/l BAP with 1.5 mg/l NAA (Table 2). In this study, we observed that increased cytokinin (BAP) over auxin (NAA) showed a better response to the percentage of shoot induction for all three okra accessions. Percent with initiate shoot increased with increasing in the concentration of BAP over the concentration of NAA in the treatments by 31.7% and 1mg/l BAP with 0.5 mg/l with NAA with 1mg/l BAP with 1.5mg/l with NAA and1.5mg/l BAP with 1.5mg/l with NAA. Growth medium without cytokinins responded to weaker shoot induction. This indicates that cytokinins are essential for shoot initiation. In agreement with the current study, the highest percentage of shoot initiation was recorded in MS medium containing 2.0 mg/l BAP (Rizwan *et al*., 2018).

Cytokinin is necessary for shoot initiation during micropropagation hormonal balance is generally believed to be a key factor in regulating morphogenesis in explants. The balance and ratios of hormones present help to influence plant reactions: they possibly regulate enzymatic reactions in the plant by amplifying them, influencing plant growth results *(Dhande et al*., 2012).

### Shoot Multiplication

From these trials, media with AC and AgNO_3_ showed better result after two weeks’ of culture. AgNO_3_ was more effective in shoot initiation and leaf formation (Figure 2).

**Figure 2:**
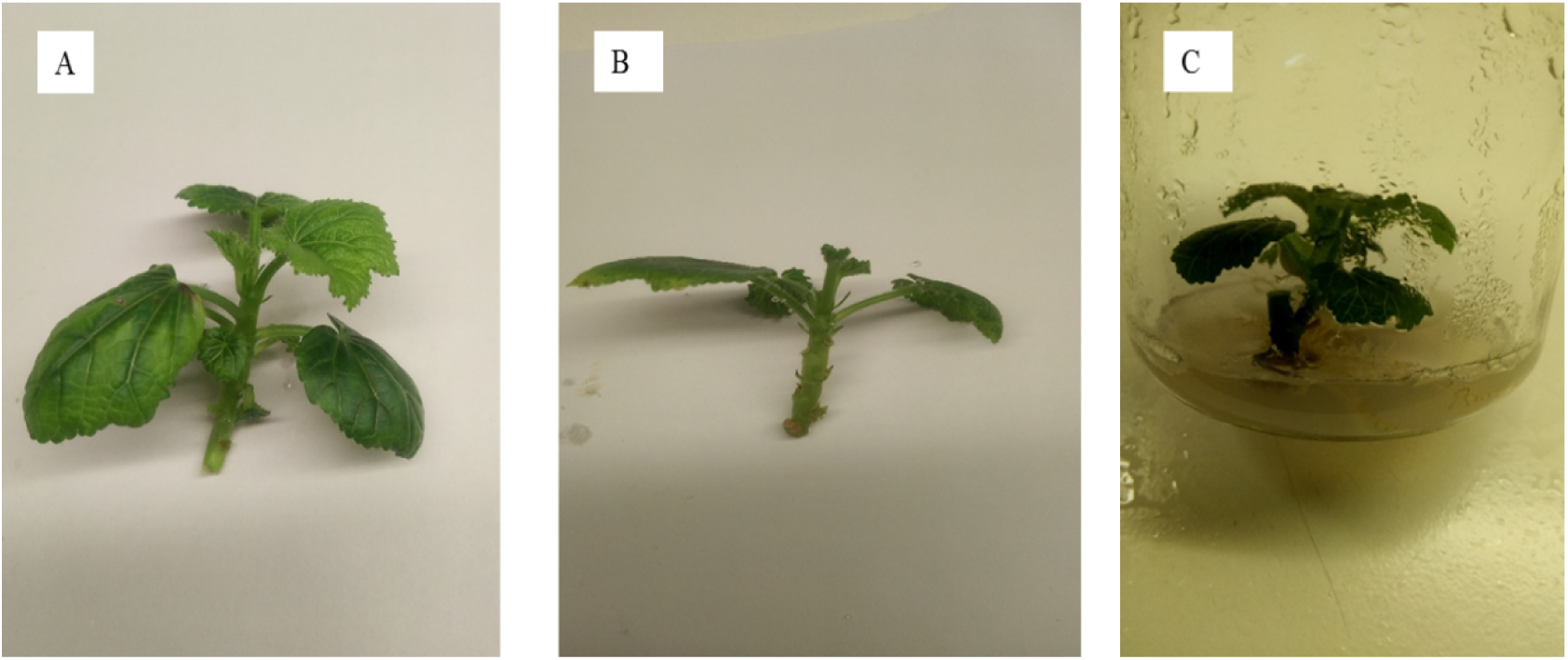
Explants on medium supplemented with PGRs and AgNO3 after two weeks’ of culture: A) Media supplemented with 2 mg/l AgNO_3_, 0.5 mg/l BAP and 0.1 mg/l NAA of Mandura accession; B) Media supplemented with 2 mg/l AgNO_3_, 0.5 mg/l BAP and 0.1 mg/l NAA of Akobo accession; C) Media supplemented with 2 mg/l AgNO_3_, 0.5 mg/l BAP and 0.1 mg/l NAA of Diga leka accession.

However, reduction in the amount of NAA and temperature and increased intensity of light were not successful in controlling the dropping of leaves. After two weeks of culture, leaves were dropped in media with reduced amounts of NAA and in media with adjusted temperature and light intensity. Accordingly, for the subsequent soot multiplication stage, only silver nitrate was used. The addition of regeneration enhancers such as silver nitrate (AgNO_3_) in the growth medium plays a vital role in shoot multiplication and is found in various economically important plants such as *Vigna mungo* (Mookkan & Andy, 2014), *Gossypium hirsutum* (Prem Kumar *et al*., 2016), *Arachis hypogaea (*Ozudogru *et al*., 2005), *Cucumis sativus* (Venkatachalam *et al*., 2018), *Catharanthus roseus* (Panigrahi *et al*., 2018) and *Prosopis cineraria* (Venkatachalam *et al*., 2017). Therefore, media supplemented with AgNO_3_ (2 mg/l) and BAP at different concentrations (0, 0.1, 0.25, 0.5, and 1 mg/l) was used for further shoot multiplication. After four weeks of explant culture, the mean values of shoot number, leaf number, and shoot length were recorded.

### Mean Number of Shoot per Explant

Statistical analysis revealed no significant difference (P>0.05) in the interaction of the factors. Similarly, there was no significant difference in the number of shoots per explant between the accessions. However, the number of shoots initiated per explant was significantly (P<0.01) influenced by BAP (Table A3).

The maximum mean number of shoots was recorded on MS media supplemented with 1.5 mg/l BAP + 2 mg/l AgNO_3_ for the three accessions followed by 1 mg/l BAP + 2 mg/l AgNO_3_. Whereas, the lowest mean number of shoots were recorded on hormone free media in all three accessions followed by 0.1 mg/l + 2 mg/l AgNO_3_ (Table 3). This study revealed that increasing the concentration of BAP is important for the production of more shoots. In hormone-free media, only one shoot was produced, which was due to cytokinin (BAP). Therefore, cytokinins are essential for the regeneration of a greater number of shoots. (Rani, 2007), obtained the same result as the present experiment, which recorded the maximum number of shoots on media supplemented with 1 mg/l BAP and the least number of multiple shoots in MS media free from growth hormones. According to the findings of (Rizwan *et al*., 2018), the maximum number of shoots recorded on the media supplemented with 1.5 mg/l BAP + 1.5 mg/1 KIN + 3.0 mg/1 AgNO_3_. In this study, a better response to the number of shoots per explant was observed with high levels of cytokinin (BAP) and AgNO_3_. This might be due to the effect of cytokinins on the regulation of cell division, which leads to shoot bud formation in the explant.

**Table 3.** Effect of BAP on number of shoot and leaf of *in vitro* initiated explants of Okra accessions during multiplication.

| BAP concentration (mg/l) | Number of shoot per explant | Number of leaf per shoot | Length of shoot of explants |
| --- | --- | --- | --- |
| 0 | 1.22±0.16<sup>d</sup> | 1.78±0.18<sup>d</sup> | 5.1±0.07<sup>a</sup> |
| 0.1 | 1.78±0.16<sup>c</sup> | 2.67±0.18<sup>c</sup> | 4.43±0.07<sup>b</sup> |
| 0.25 | 2.33±0.16<sup>b</sup> | 3.22±0.18<sup>b</sup> | 4.2±0.07<sup>c</sup> |
| 0.5 | 2.67±0.16<sup>b</sup> | 3.56±0.18<sup>ab</sup> | 3.4±0.07<sup>d</sup> |
| 1 | 3.22±0.16<sup>a</sup> | 3.78±0.18<sup>a</sup> | 3.04±0.07<sup>e</sup> |
| 1.5 | 3.22±0.16<sup>a</sup> | 4±0.18<sup>a</sup> | 2.4±0.07<sup>f</sup> |
| LSD (5%) | 0.47 | 0.52 | 0.24 |
| CV (%) | 19.68 | 17.2 | 6.01 |

CV, coefficient of variation; LSD= Least Significant Difference; SE= Standard Error; BAP, 6-benzylaminopurine. Means with different letters in a column are statistically significant at $p < 0.05$ . Values are presented as the mean ± SE.

### Mean Number of Leaf per Shoot

The number of leaves per shoot in *in vitro* nodal culture of the three accessions of Okra varied among different concentrations and combinations of plant growth regulators. Statistical analysis revealed no significant difference (P>0.05) in the interaction of the factors. Similarly, there was no significant difference in the number of leaves per explant between the accessions. The number of leaves initiated per explant was significantly (P<0.01) influenced by the main effect of BAP.

The maximum number of leaves per shoot was recorded on MS media supplemented with 1.5 mg/l BAP +2 mg/l AgNO_3_ followed by 1 mg/l BAP +2 mg/l AgNO_3_ for all three accessions. The lowest mean number of leaves per shoot was recorded in the hormone-free media (Table 3). According to (Prem Kumar *et al*., 2016) the maximum number of shoots was recorded on media supplemented with 1 mg/l BAP and silver nitrate. In the present study, increasing concentrations of BAP were essential for inducing the maximum number of leaves per initiated shoot. In this study, the number of shoots and leaves increased with increasing concentrations of BAP (Figure 3). Therefore, the current finding is in line with the result by (Wondye Ayaliew et al., 2026) who recorded the highest mean number of shots (4.66 ± 0.32) through 1.5 mg/l of BAP.

**Figure 3:**
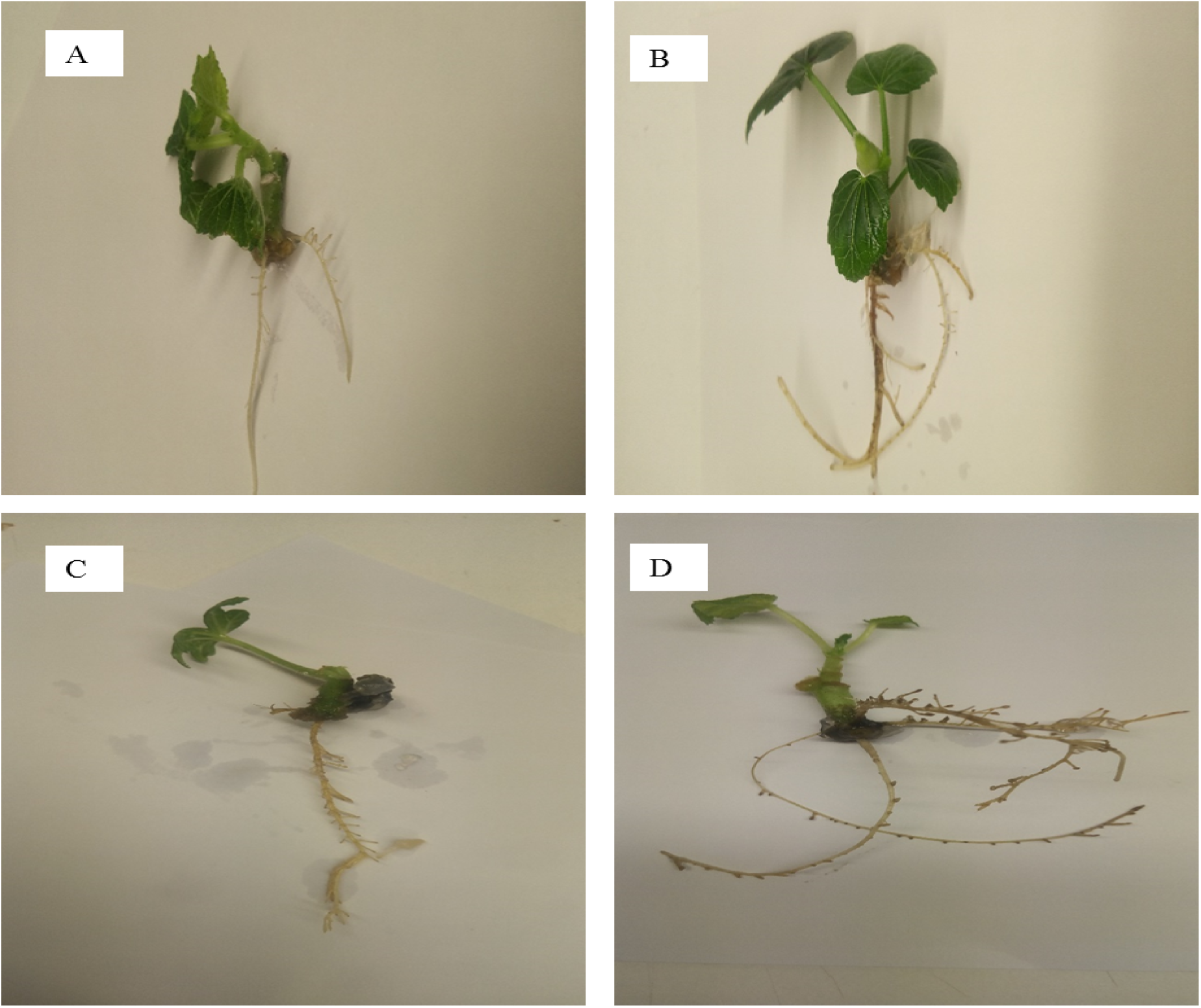
Roots induced from regenerated shoots. A) Media supplemented with 0.5 mg/l IBA of Akobo accession. B) Media supplemented with 0.25 mg/l of IBA Diga leka accession. C) Media without IBA of Diga leka accession. D) Media supplemented with 2mg/l of IBA of accession Mandura.

### Mean Shoot Length

Statistical analysis revealed that there was no significant difference (P>0.05) in the interaction between the two factors. Similarly, there was no significant difference in the length of shoots per explant between the accessions. Shoot length per explant was significantly (P<0.01) influenced by the main effect of BAP. (Table A3).

The tallest shoots of the explants were recorded on MS medium without plant growth regulators (BAP) for all accessions. Whereas, the shortest length of the explant was recorded on the MS media supplemented with 1.5 mg/l of BAP + 2 mg/l AgNO_3_ followed 1 mg/l of BAP with 2 mg/l AgNO_3_ for three accessions of Okra (Table 3). In this study, better results were obtained for shoot length per explant on MS media free from plant growth regulators for all accessions. This may be due to the lower number of shoots produced in the media. The results of this study also showed that shoot length decreased with increasing concentrations of BAP (Table 3). The tallest shoot was recorded on the media supplemented with 1.5 mg/l BAP + 1.5 mg/1 KIN + 3.0 mg/1 AgNO_3_ (Rizwan *et al*., 2018).

### Root Induction

After multiplication, well-developed shoots were transferred to rooting media. Auxins are mainly used for root induction, and their effects vary depending on the type and concentration used in different plant species (Swamy *et al*., 2002). In this study, IBA at different concentrations was used to induce root growth. IBA has yielded good results for root number because it is very effective in increasing endogenous auxin content and shows higher stability against catabolism and activation by conjugation with growth inhibitors (Hasan *et al*., 2010).

### Days to Root Induced

Statistical analysis revealed no significant difference (P>0.05) in the interaction of the factors. The number of days required for root induction was significantly (P<0.01) and significantly (P<0.05) influenced by the main effects of growth regulators and accessions, respectively (Table A4).

The minimum mean number of days to root initiation was recorded for explants cultured on MS medium supplemented with 0.5 mg/l IBA followed by 1 mg/l IBA. The maximum mean number of days to root initiation was recorded for explants cultured on media without plant growth regulators, followed by 2 mg/l IBA (Table 4). The results of the present study revealed that the number of days to initiate root increased with increasing in IBA concentration in the treatments by 35.8% from 0.5 mg/l of IBA to 2 mg/l of IBA. The accession Mandura also took fewer days to root induction than accessions Akobo and Diga Leka, which took more days to induce roots. The current finding is in accordance with a previous report on Okra where IBA alone was superior for induction of root elongation, as reported by (Kabir *et al*., 2016) writen that the root growth after transferring elongated shoots to the rooting medium. The differences in rooting responses may be a result of genotype or cultural conditions. Similar to the current study, root induction was recorded between 10-15 days Cotton crops grown in media supplemented with 4.92 μM IBA (Pathi & Tuteja, 2013).

**Table 4.** Effect of IBA on days and percent of root induced and on number of roots per explant of *in vitro* initiated shoots of the three Okra accessions.

| IBA concentration (mg/l) | DRI | Number of root per explant | Length of root per explant |
| --- | --- | --- | --- |
| 0 | 11.78±0.24<sup>a</sup> | 1.22±0.17<sup>d</sup> | 9.5±0.37<sup>a</sup> |
| 0.25 | 8.33±0.24<sup>d</sup> | 2±0.17<sup>c</sup> | 9.01±0.37<sup>ab</sup> |
| 0.5 | 6.56±0.24<sup>f</sup> | 2.56±0.17<sup>b</sup> | 8.93±0.37<sup>ab</sup> |
| 1 | 7.33±0.24<sup>e</sup> | 2.89±0.17<sup>b</sup> | 7.83±0.37<sup>c</sup> |
| 1.5 | 9.22±0.24<sup>c</sup> | 3.44±0.17<sup>a</sup> | 8.38±0.37<sup>bc</sup> |
| 2 | 10.22±0.24<sup>b</sup> | 3.78±0.17<sup>a</sup> | 9.94±0.37<sup>a</sup> |
| LSD (5%) | 0.69 | 0.49 | 1.18 |
| Accessions |  |  |  |
| Akobo | 8.9±0.17<sup>ab</sup> | 2.44±0.12<sup>b</sup> | 8.34±0.26<sup>b</sup> |
| Diga leka | 9.28±0.17<sup>a</sup> | 2.44±0.12<sup>b</sup> | 8.68±0.26<sup>b</sup> |
| Mandura | 8.56±0.17<sup>b</sup> | 3.06±0.12<sup>a</sup> | 9.78±0.26<sup>a</sup> |
| LSD (5%) | 0.49 | 0.34 | 0.83 |
| CV (%) | 8.08 | 19.23 | 12.53 |

CV, coefficient of variation; LSD= Least Significant Difference; SE= Standard Error; IBA= Indole-3-butyric acid; DRI= Days to root induction Means with different letters in a column are statistically significant at $p < 0.05$ . Values are presented as the mean ± SE.

### Percent of Root Induced

Statistical analysis showed that there was no significant (P>0.05) difference in the percent of root induced per shoot among the main effects of plant growth regulators and accessions. MS media supplemented with a plant growth regulator (IBA) and media without a growth regulator (IBA) showed the same response (Table A4). However, there was a difference in the number of roots and the length of roots induced per shoot.

### Mean Number of Root Induced per Shoot

The number of roots induced per shoot in *in vitro* nodal culture of the three accessions of Okra varied among different concentrations and combinations of plant growth regulators. Statistical analysis showed that there was no significant (P>0.05) difference in the number of roots per initiated shoot among the interaction of the main effect of accessions and plant growth regulator (IBA). However, there was a highly significant (P<0.01) difference in the number of roots per shoot among the main effects of plant growth regulator and accessions.

The maximum mean number of roots per shoot was recorded on MS media supplemented with 2 mg/l IBA followed by 1.5 mg/l IBA and 1 mg/l IBA. The lowest mean number of roots per shoot was recorded in media without growth regulators (Table 4). The result of the present study revealed that the number of root induced for all accessions were increased with increasing in the concentration of IBA in the treatments by 47.1% from 0.25 mg/l of IBA to 2 mg/l of IBA. In the accessions, the maximum mean number of roots was recorded under accession Mandura, followed by accessions Akobo and Diga Leka. According to (Rizwan *et al*., 2018), the maximum number of roots per shoot was obtained in media supplemented with 1 mg/l mg/L IBA. (Patil *et al*., 2017) also observed similar result in which maximum number of root induction on half strength media supplemented with 1.5 mg/l IBA in *Abutilon ranadei*. (Manokari *et al*., 2016) also observed the maximum number of roots in media supplemented with 2 mg/l IBA in *Hibiscus subdariffa*. In this study, we observed that increasing the concentration of auxin (IBA) was important for the induction of more roots (Figure 4).

**Figure 4:**
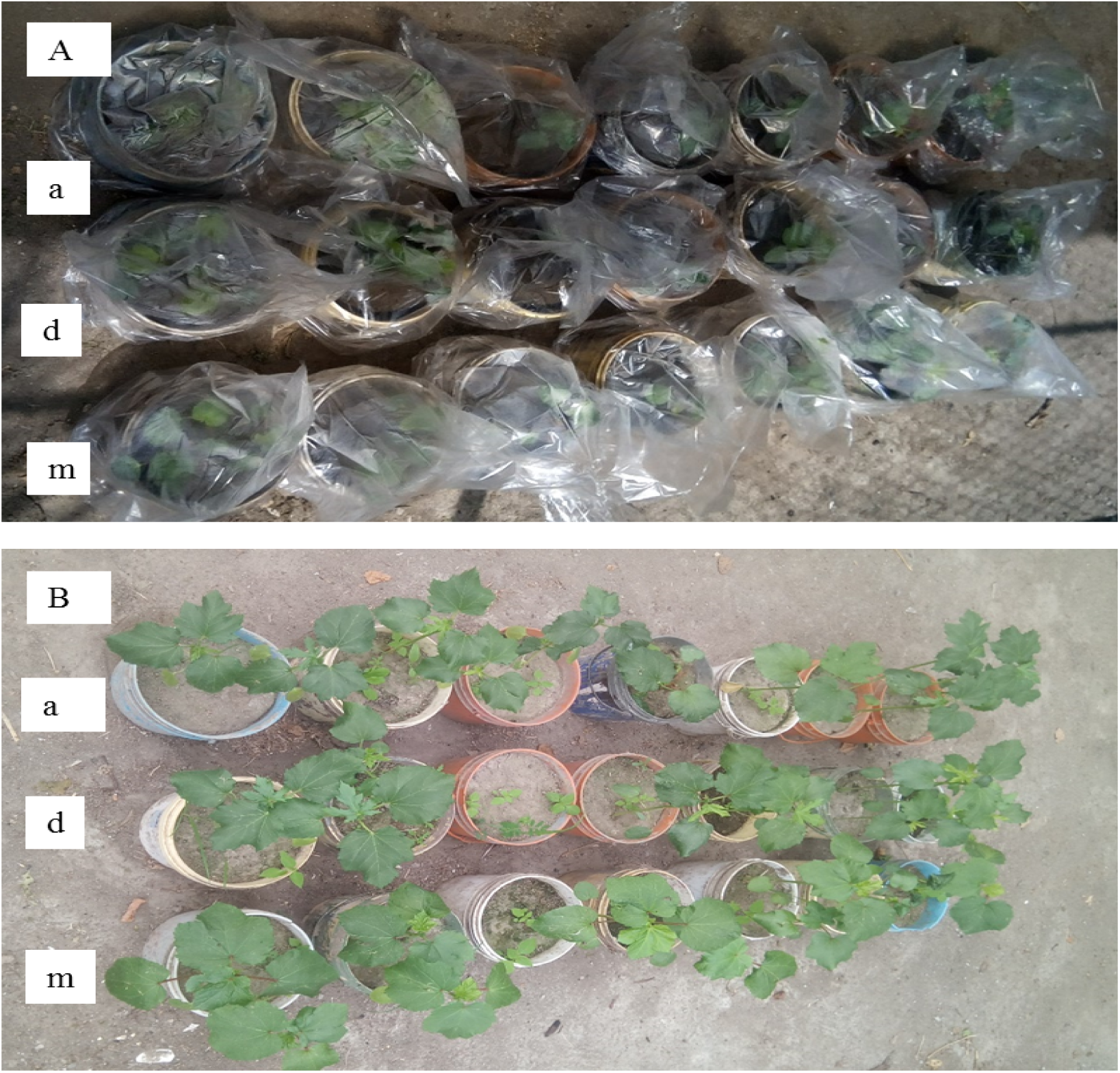
Acclimatized plantlets of Okra accessions in the net house. A) Plantlets covered by plastic bags for the first 15 days; B) After 30 days of growth in the net house. a=Akobo accession; d=Diga leka accession; and m=Mandura accession.

### Mean Length of Root Induced per Shoot

Statistical analysis showed, there was no significant difference (P>0.05) between the interactions of the main effect. However, there was a significant (P<0.01) influence on the length of roots per shoot among the main effects of plant growth regulator and accessions.

The maximum mean root length was obtained on media supplemented with 2 mg/l, IBA followed by media free of plant growth regulators. The lowest root length in the experiment was recorded on media supplemented with 1 mg/l IBA followed by 1.5 mg/l IBA. The maximum mean root length of accession was obtained from accession Mandura, followed by accessions Diga leka and Akobo (Table 4). This result showed that the length of the roots varied with the concentration IBA at high and low levels. (Patil *et al*., 2017) also observed similar result in which maximum length of root on media supplemented with 1.5 mg/l and 1 mg/l IBA in *Abutilon ranadei*. Similarly, different researchers have reported the effect of different forms of auxin on root length. (Manokari *et al*., 2016) observed maximum root length in media supplemented with 2 mg/l IBA in *Hibiscus subdariffa*. (Dhande *et al*., 2012) also obtained the longest roots on media supplemented with 1 mg/l IAA. According to (Wondye Ayaliew et al., 2026) mean length of the roots was increased up to treatment (2 mg/l) of NAA in a linear fashion, while it was reduced above (2.5 mg/l), which might indicate the toxic effect of NAA above 2 mg/l concentration.

### Acclimatization

In the present study, well-rooted plantlets were rinsed in water to remove traces of medium with agar and sucrose and grown in a 2:1:1 mixture of soil, compost, and sand in plastic pots. As the regenerated plantlets needed gradual acclimatization for their survival in the field condition after growing in the controlled environment instead of transferring directly to the field, plantlets were left for 4 days in the greenhouse at controlled temperature and optimum relative humidity. After four days, all acclimatized plants were transferred to a net house and the pots were covered with plastic bags with holes for aeration (Figure 5) for hardening. After 30 days hardening of the plantlets, 73.33 % of Mandura accession and 66.67 of accession Diga Leka and Akobo were survived (Table 5). (Rani, 2007) observed that only 25 percent of germinated plants survived after one month of acclimatization. The loss of some plantlets may be due to differences in the genotype in adaptation to the new environment (*ex vitro*). The reduced development of cuticle under *in vitro* conditions and the drop in relative humidity from nearly 100% in the cuticle vessels to much lower values in the greenhouse may result in excessive water loss and death (Biradar *et al*., 2010). There were no observable variations in morphological and growth characteristics between *ex vitro* and *in vitro* raised plants in the pots.

**Table 5.** Survival rate (%) of plantlets transferred from *in vitro* culture to outdoor for acclimatization of the three accessions of Okra.

| Accessions | Number of plantlets transferred to acclimatization | Number of plantlets survived | Number of plantlets died | Percentage of survived plantlets | Percentage of died plantlets |
| --- | --- | --- | --- | --- | --- |
| Akobo | 15 | 10 | 5 | 66.67 | 33.33 |
| Diga leka | 15 | 10 | 5 | 66.67 | 33.33 |
| Mandura | 15 | 11 | 4 | 73.33 | 26.67 |

## SUMMARY AND CONCLUSION

Okra (*Abelmoschus esculentus* L. Moench) is an economically important vegetable. Okra also possesses many medicinal and nutritional properties. Owing to its many advantages, the propagation and production of crops is of paramount importance. However, conventional propagation methods are not adequate for efficient production of this crop. Plant tissue culture is the best alternative for improving okra production. However, *in vitro* propagation study of okra crops in Ethiopia has not yet been conducted. Therefore, this study was designed to develop a protocol for *in vitro* regeneration of Okra of Ethiopian accessions, namely, Akobo, Diga leka, and Mandura accessions through nodal cutting.

To culture nodal cuttings, the mother plant was planted in a net house, and the explant was taken after a month of cultivation. The explant was sterilized and shoot initiation was performed. Shoot initiation in tissue culture is controlled by *in vitro* media composition. Hormonal balance plays a major role in controlling the growth of tissues. The ratio of auxin to cytokinin determines the response of *in vitro* cultured tissues. A higher cytokinin to auxin ratio promotes shooting, while the reverse promotes rooting, and the balanced levels of the two hormones promote callus formation.

In this study, different hormone concentrations and combinations were used for shoot initiation. From shoot inducing media the minimum days and highest percentage of shoot induction from nodal explant was obtained from MS medium containing 1 mg/l BAP and 0.5 mg/l NAA for all three accessions.

For the shoot multiplication stage, different concentrations of BAP and 2 mg/l mg/L AgNO_3_ were used for all three accessions. At this stage, MS media supplemented with 1.5 mg/l of BAP and 2 mg/l of AgNO_3_ was necessary for maximum shoot and leaf number for all three accessions. On the other hand, the longest shoot was obtained on MS medium free from plant growth regulators and AgNO_3_. This shows that the number of shoots and the length of shoots are inversely proportional. If the number of shoots increases, the length decreases, and the reverse is true.

The rooting stage was followed by the shoot multiplication stage. For this stage, IBA at different concentrations was used to induce roots through the initiated shoots. In this study, the minimum number of days for root induction was recorded on MS medium supplemented with 0.5 mg/l IBA. The percentage of root induction was not significant because of the main effects and their interaction. This implies that in all treatments, including media free from growth hormone-induced roots. The maximum mean number of roots was recorded on MS medium supplemented with 2 mg/l IBA. However, the highest root length was recorded in MS medium free from plant growth regulators.

The last stage of the experiment involved acclimatization of the plantlets *ex vitro*. For acclimatization, plantlets from nodal culture experiments were transferred to net house environment. After a month of acclimatization maximum percentage of survival (73.33%) was obtained from explants regenerated from Mandura accessions. Therefore, it can be concluded that okra can be regenerated in tissue culture and established outside. Finally, in this study, an efficient regeneration protocol from nodal culture experiments was developed for large-scale multiplication, propagation, and conservation of okra. This technique can be used for the subsequent mass propagation of true-to-type plants and genetic transformation of this very important vegetable crop.

Future perspectives based on the present study should focus on the following areas.

✓ *In vitro* regeneration of okra is possible. Therefore, further research on improving the productivity of okra, such as genetic transformation, should be conducted.
✓ Explants other than node cultures such as leaves, shoot tips, and roots for *in vitro* propagation of okra should be considered to investigate whether they can offer better and more efficient responses.
✓ Different strength of MS media should be tested for rooting.
✓ Indirect regeneration (through callus) of plants should be studied in different accessions.
✓ Further research on the leaf-dropping problem should be conducted. AgNO_3_ at different concentration should also be studied.
✓ Different types of cytokinins for shoot multiplication and auxins in root formation should be tested.

## Acknowledgment

Samuel Arega Asfaw is thankful to his advisor, Dr. Meseret Tesema, for her support and fruitful suggestions and advice from proposal development to the completion of the research work. The author SAA has benefited and learned a lot from discussions on various aspects of the research. He is also highly indebted to his co-advisor, Dr. Zerihun Demerew, for his essential role in guiding this research work. The author is also grateful to Assosa University for sponsoring MSc opportunities. SAA performed experiments as part of his master’s thesis. All studies were carried out under the supervision of Dr. Mseret Tesema.

## Authors contributions

**Samuel Arega Asfaw**: Conceptualization, Data curation, analysis, material collection Methodology, Resources, Software, Supervision, Validation, Visualization, Writing – original draft, writing – review and editing; **Meseret Tesema** Conceptualization, Data curation, Data analysis, Supervision, Investigation, Resources, Software, Writing review and editing; **Zerihun Demerew:** Conceptualization, Formal analysis, Methodology, Software, Validation, Writing – original draft; **Sualih Gobezie**: Conceptualization, Data curation, Data analysis, Software, Validation, Visualization, Writing – review and editing; **Wondye Ayaliew Shiferaw**: Conceptualization, Methodology, Software, Validation, Visualization, Writing – original draft, review and editing.

## Ethical approval

Not applicable, hence this study did not involve human participants or animals.

## Data availability statement

The data that support the findings of this study are available in Zenodo at https://doi.org/10.5281/zenodo.21155681. These data’s (Excel data, Figure data and Table data) derived from the following resources available in the public domain.

## APPENDIX

**Appendix table 1:**
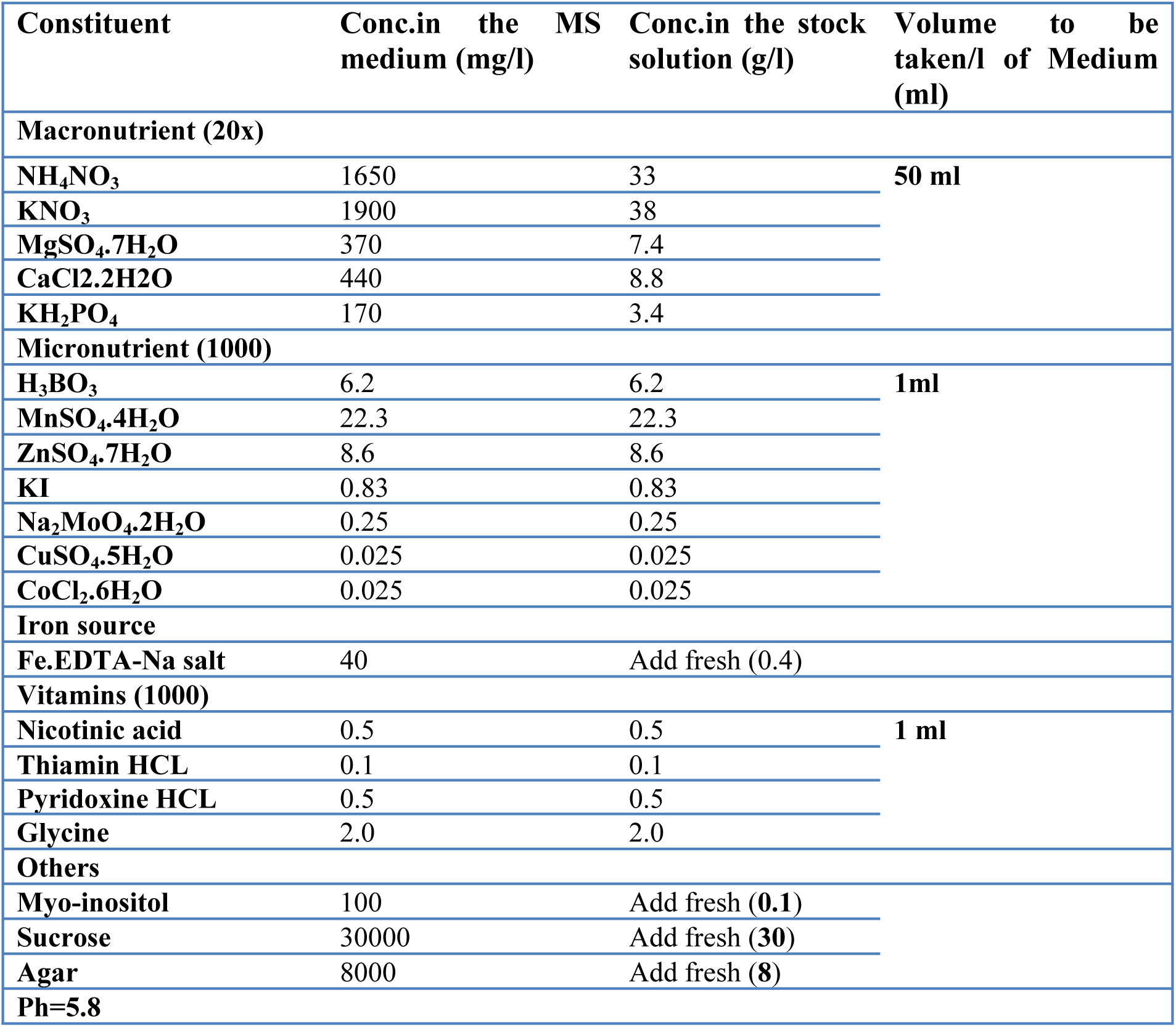
List of MS media components and their concentrations in stock solutions and media.

**Appendix table 2:** Mean square estimates of in vitro shoot regeneration and growth parameters of the three okra accessions as affected by the PGRs (BAP and NAA) combination.

| Source of variation | DF | Percent of shoot initiation |  | Days to shoot initiation |  |
| --- | --- | --- | --- | --- | --- |
|  |  | MS | F | MS | F |
| PGR | 9 | 0.2 | 21.81** | 35.67 | 138.22** |
| Accession | 2 | 0.001 | 0.09<sup>ns</sup> | 0.45 | 1.75<sup>ns</sup> |
| PGR*Accession | 18 | 0.008 | 0.75<sup>ns</sup> | 0.23 | 0.89<sup>ns</sup> |
| Error | 120 | 0.01 |  | 0.26 |  |
| CV (%) |  | 15.45 |  | 6.05 |  |

\*\* = significant at 1 % probability levels; <sup>ns</sup>= non-significant at 5 %; DF = Degree of freedom; PGR= Plant growth regulator; MS=mean squares; F=F value of ANOVA and CV= coefficient of variation.

**Appendix table 3:** Mean square estimates of in vitro shoot multiplication and growth parameters of the three okra accessions affected by BAP.

| Source of variation | DF | Number of shoot |  | Shoot length |  | Number of leaf |  |
| --- | --- | --- | --- | --- | --- | --- | --- |
|  |  | MS | F | MS | F | MS | F |
| BAP | 5 | 5.76 | 23.94** | 5.81 | 140.74** | 6.12 | 20.66** |
| Accession | 2 | 0.46 | 1.92<sup>ns</sup> | 0.006 | 0.09<sup>ns</sup> | 0.39 | 1.31<sup>ns</sup> |
| BAP*Accession | 10 | 0.26 | 1.09<sup>ns</sup> | 0.01 | 0.18<sup>ns</sup> | 1.84 | 1.84<sup>ns</sup> |
| Error | 45 | 0.22 |  | 0.06 |  | 1.24 |  |
| CV (%) |  | 19.68 |  | 6.01 |  | 17.2 |  |

\*\* = significant at 1 % probability levels; <sup>ns</sup>= non-significant at 5 %; DF = Degree of freedom; BAP= 6-benzylaminopurine; MS=mean squares; F=F value of ANOVA and CV= coefficient of variation.

**Appendix table 4:**
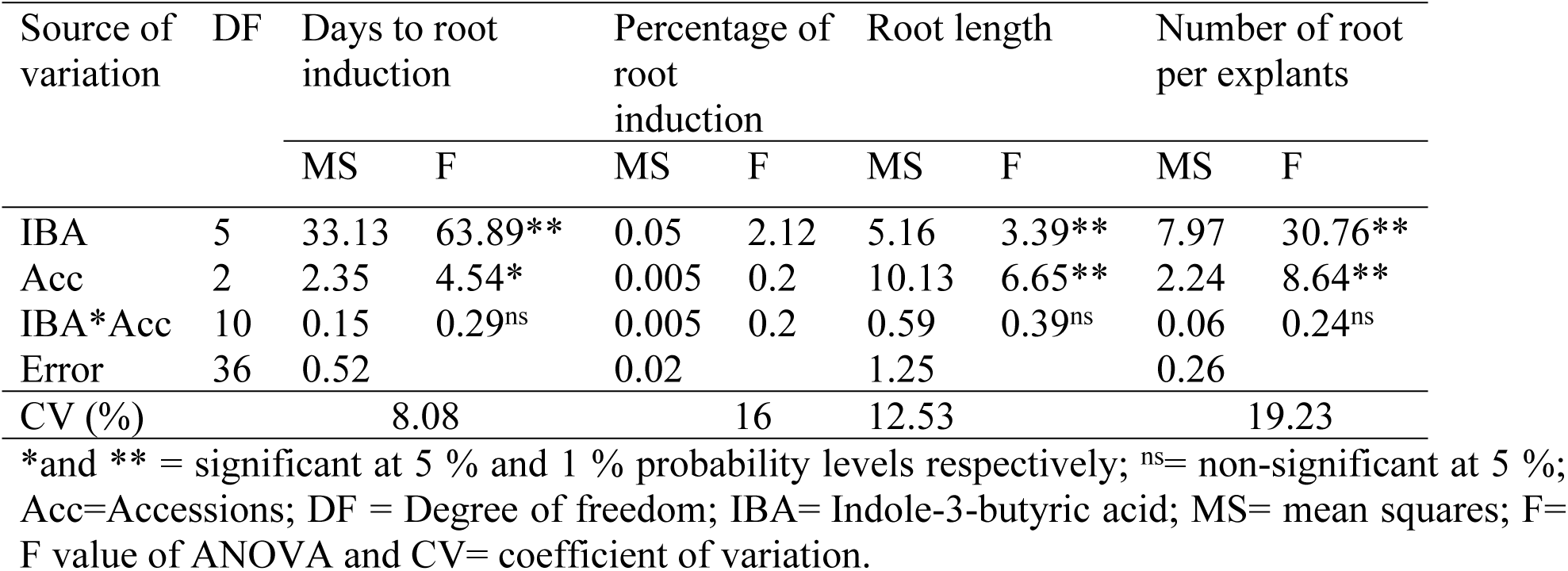
Mean square estimates of in vitro rooting and growth parameters of the three okra accessions affected by IBA.

## REFERENCES

Akinyele, B., & Temikotan, T. (2007). Effect of variation in soil texture on the vegetative and pod characteristics of okra (Abelmoschus esculentus (L.) Moench). International Journal of Agricultural Research, 2(2), 165–169.

Arya, I., Chauhan, S. S. S., & Arya, S. (2009). Micropropagation of superior eucalyptus hybrids FRI-5 (Eucalyptus camaldulensis Dehn x E. tereticornis Sm) and FRI-14 (Eucalyptus torelliana FV Muell x E. citriodora Hook): A commercial multiplication and field evaluation. African Journal of Biotechnology, 8(21), 5718.

Biradar, S., Biradar, D., Patil, V., Patil, S., & Kambar, N. (2010). In vitro plant regeneration using shoot tip culture in commercial cultivar of sugarcane. Karnataka Journal of Agricultural Sciences, 22(1).

Blackmon, W., Reynolds, B., & Postek, C. (1981). Production of somatic embryos from callused cantaloupe hypocotyl explants. HortScience,

Dabire-Binso, C. L., Ba, M. N., Some, K., & Sanon, A. (2009). Preliminary studies on incidence of insect pest on okra, Abelmoschus esculentus (L.) Moench in central Burkina Faso. African Journal of Agricultural Research, 4(12), 1488–1492.

Dhande, G., Patil, V. M., Raut, R., Rajput, J., & Ingle, A. (2012). Regeneration of okra (Abelmoschus esculentus L.) via apical shoot culture system. African Journal of Biotechnology, 11(86), 15226–15230.

El-Kader, A., Shaaban, S., & El-Fattah, M. (2010). Effect of irrigation levels and organic compost on okra plants (Abelmoschus esculentus L.) grown in sandy calcareous soil. Agriculture and Biology Journal of North America, 1(3), 225–231.

FAOSTAT. (2016). countries by commodity.

Feyissa, T., Welander, M., & Negash, L. (2005). In vitro regeneration of Hagenia abyssinica (Bruce) JF Gmel.(Rosaceae) from leaf explants. Plant Cell Reports, 24(7), 392–400.

Haider, S., Islam, R., Kamal, A., Rahman, S., & Joarder, O. (1993). Direct and indirect organogenesis in cultured hypocotyl explants of Abelmoschus esculentus (L.) moench [in Bangladesh]. Plant Tissue Culture, 3.

Hasan, S. Z. U., Ahmad, T., Hafiz, I. A., & Hussain, A. (2010). Direct plant regeneration from leaves of Prunus rootstock GF677 (Prunus amygdalus× P. persica). Pak. J. Bot, 42(6), 3817–3830.

Kabir, M., Ahmed, S., & Akhond, M. (2016). Organogenesis in okra (Abelmoschus esculentus L. Moench.): A plant recalcitrant to tissue culture. Bangladesh Journal of Agricultural Research, 41(3), 521–528.

Kalimuthu, K., Paulsamy, S., Senthilkumar, R., & Sathya, M. (2007). In vitro propagation of the biodiesel plant Jatropha curcas L. Plant Tissue Culture and Biotechnology, 17(2), 137–147.

Kirakosyan, A., Vardapetyan, R., Charchoglyan, A., Yamamoto, H., Hayashi, H., & Inoue, K. (2001). The Effect of Cork Pieces on Pseudohypericin Production in Cells ofHypericum perforatumShoots. Russian Journal of Plant Physiology, 48(6), 816–819.

Kucharek, T. (2004). Florida plant disease management guide: Okra. Plant Pathology Department document PDMG-V3-41. Florida Cooperative Extension Service, Institute of Food and Agricultural Sciences, *University of Florida*, *Gainesville*, *FL*, 50-55.

Lengsfeld, C., Titgemeyer, F., Faller, G., & Hensel, A. (2004). Glycosylated compounds from okra inhibit adhesion of Helicobacter pylori to human gastric mucosa. Journal of Agricultural and Food Chemistry, 52(6), 1495–1503.

Mangat, B., & Roy, M. (1986). Tissue culture and plant regeneration of okra (Abelmoshus esculentus). Plant Science, 47(1), 57–61.

Manokari, M., Ravindran, C., & Shekhawat, M. S. (2016). Optimization of in vitro regeneration in Hibiscus subdariffa L.–An important medicinal plant. World Scientific News(34), 51–62.

Mookkan, M., & Andy, G. (2014). AgNO3 boosted high-frequency shoot regeneration in Vigna mungo (L.) Hepper. Plant Signaling & Behavior, 9(10), e972284.

Murashige, T., & Skoog, F. (1962). A revised medium for rapid growth and bio assays with tobacco tissue cultures. Physiologia plantarum, 15(3).

NMA. (2012). National meteorological agency, Hawassa branch directorate, Southern Nations, Nationalities and Peoples Regional State, Ethiopia.

Ozudogru, E. A., Ozden-Tokatli, Y., & Akcin, A. (2005). Effect of silver nitrate on multiple shoot formation of Virginia-type peanut through shoot tip culture. In Vitro Cellular & Developmental Biology-Plant, 41(2), 151–156.

Panigrahi, J., Dholu, P., Shah, T. J., & Gantait, S. (2018). Silver nitrate-induced in vitro shoot multiplication and precocious flowering in Catharanthus roseus (L.) G. Don, a rich source of terpenoid indole alkaloids. Plant Cell, Tissue and Organ Culture (PCTOC), 132(3), 579–584.

Pathi, K. M., & Tuteja, N. (2013). High-frequency regeneration via multiple shoot induction of an elite recalcitrant cotton (Gossypium hirsutum L. cv Narashima) by using embryo apex. Plant Signaling & Behavior, 8(1), e22763.

Patil, S., Mane, J., Umdale, S., & Chavan, J. (2017). Direct and indirect shoot organogenesis strategies for propagation and conservation of Abutilon ranadei Wooder & Stapf-a critically endangered plant. Adv Cel Sci & Tissue Cul *2017;* 1 (1): 1–4*. 2 Adv Cel Sci & Tissue Cul 2017 Volume 1 Issue*, *1*, 3.

Prem Kumar, G., Sivakumar, S., Siva, G., Vigneswaran, M., Senthil Kumar, T., & Jayabalan, N. (2016). Silver nitrate promotes high-frequency multiple shoot regeneration in cotton (Gossypium hirsutum L.) by inhibiting ethylene production and phenolic secretion. In Vitro Cellular & Developmental Biology-Plant, 52(4), 408–418.

Rani, M. R. (2007). Standardization of Protocol for Regeneration and Genetic Transformation of Okra (Abelmoschus Esculentus [L.] Moench) ANGRAU College of Agriculture: Agricultural Biotechnology].

Rizwan, H., Irshad, M., He, B., Liu, S., Debnath, B., Mitra, S., Li, M., Lu, X., Sun, Y., & Qiu, D. (2018). SILVER NITRATE (AgNO 3) BOOSTED HIGH-FREQUENCY MULTIPLE SHOOT REGENERATION FROM COTYLEDONARY NODE EXPLANTS OF OKRA (ABELMOSCHUS ESCULENTUS L.). Applied Ecology & Environmental Research, 16(3).

Roy, A., Shrivastava, S. L., & Mandal, S. M. (2014). Functional properties of Okra Abelmoschus esculentus L.(Moench): traditional claims and scientific evidences. Plant science today, 1(3), 121–130.

Santarém, E. R., & Astarita, L. V. (2003). Multiple shoot formation in Hypericum perforatum L. and hypericin production. Brazilian journal of plant physiology, 15(1), 43–47.

SAS (Ed.). (2002). (Statistical Analysis System). SAS/STAT. Guide version 9.0. SAS, Institute Inc. Cary North Carolina, USA.

Schippers, R. R. (2000). African indigenous vegetables: an overview of the cultivated species. Natural Resources Institute/ACP-EU Technical Centre for Agricultural and ….

Swamy, S., Puri, S., & Singh, A. (2002). Effect of auxins (IBA and NAA) and season on rooting of juvenile and mature hardwood cuttings of Robinia pseudoacacia and Grewia optiva. New Forests, 23(2), 143–157.

Trigiano, R. N., & Gray, D. J. (2016). Plant tissue culture, development, and biotechnology. CRC Press.

Venkatachalam, P., Jinu, U., Gomathi, M., Mahendran, D., Ahmad, N., Geetha, N., & Sahi, S. V. (2017). Role of silver nitrate in plant regeneration from cotyledonary nodal segment explants of Prosopis cineraria (L.) Druce.: A recalcitrant medicinal leguminous tree. Biocatalysis and Agricultural Biotechnology, 12, 286–291.

Venkatachalam, P., Jinu, U., Sangeetha, P., Geetha, N., & Sahi, S. V. (2018). High frequency plant regeneration from cotyledonary node explants of Cucumis sativus L. cultivar ‘Green Long’via adventitious shoot organogenesis and assessment of genetic fidelity by RAPD-PCR technology. 3 Biotech, 8(1), 60.

Wondye Ayaliew, S., Gobena Roro, A., Demerew, Z., Arega, S., Berihun, B., & Tesema, A. A. (2026). In vitro propagation and microtuberization of yam (Dioscorea species) Hatiye landrace. Cogent Food & Agriculture, 12(1), 2631255.

Yildiz, M. (2012). The prerequisite of the success in plant tissue culture: high frequency shoot regeneration. In Recent advances in plant in vitro culture. IntechOpen.

Yonas, M., Garedew, W., & Debela, A. (2014). Multivariate analysis among okra (Abelmoschus esculentus (l.) moench) collection in South Western Ethiopia. Journal of Plant Sciences, 9(2), 43.

